# Defining transcription factor nucleosome binding with Pioneer-seq

**DOI:** 10.1101/2022.11.11.516133

**Authors:** Maria Tsompana, Patrick D. Wilson, Vijaya Murugaiyan, Christopher R. Handelmann, Michael J. Buck

**Affiliations:** Department of Biochemistry, Jacobs School of Medicine and Biomedical Sciences, State University of New York at Buffalo, Buffalo, NY, United States; Department of Biomedical Informatics, Jacobs School of Medicine and Biomedical Sciences, State University of New York at Buffalo, Buffalo, NY, United States

**Author notes:** Co-first authors (Maria Tsompana, Patrick D. Wilson).

## Abstract

Gene expression requires the targeting of transcription factors (TFs) to regulatory sequences often occluded within nucleosomes. To comprehensively examine TF nucleosome binding, we developed Pioneer-Seq. In Pioneer-seq a library of nucleosomes containing thousands of DNA sequences with TF binding sites in all possible nucleosome orientations is bound to a TF and the protein-nucleosome complex is isolated and quantified. To demonstrate Pioneer-seq we examined nucleosome binding by OCT4, SOX2, KLF4, and c-MYC. Our results demonstrate that KLF4 and SOX2 can bind close to the nucleosome dyad with nucleosome sequence being the major factor regulating TF binding across all studied TFs.

## Introduction

The interactions between proteins and chromosomal DNA underlie basic nuclear processes, such as transcriptional regulation, DNA replication, repair, and recombination, chromosome segregation, and epigenetic inheritance, as well as many fundamental biological responses, including cell growth, division, and differentiation, embryonic development, environmental stress responses, apoptosis, and disease state development. According to the human protein atlas, 8,887 proteins localize to the nucleus, with an estimated 1,500 DNA-binding transcription factors (TFs) [1]. Most TFs preferentially bind nucleosome-free DNA, which appears to be a conundrum during development because many gene regulatory regions are in nucleosomal DNA. However, there are a few TFs that belong to a specific class, known as pioneer factors, that can bind to closed chromatin and open nucleosomal domains [2–6].

Pioneer factors are not able to bind all their targets throughout the genome, indicating that there are some constraints to their binding abilities. There is evidence that the location of the TF binding site (TFBS) within a nucleosome (known as the translational setting) is one determinant of TF-binding abilities [6]. For example, binding of the glucocorticoid receptor at its TFBS near the nucleosome edge is 4-fold greater than at an identical site positioned 20 bp from the nucleosome dyad [7]. The translational setting can inhibit TF binding 2-to 100-fold [8–11]. TF binding is also influenced by the orientation of a TFBS on a nucleosome (known as the rotational setting) that results from the twist of the helical DNA structure. FoxA binds to its well-defined target site in the *Alb* enhancer (5–15 bp from the nucleosome dyad in liver cells) [12] only at specific rotational settings [13]. The ways by which TFs recognize their TFBSs (e.g., partial motif recognition) also likely influences their ability to bind nucleosomal DNA [14].

Until recently, the only way to determine whether and how a TF binds its target within nucleosomal DNA was by a low-throughput molecular biology assay, in which *in vitro-*formed nucleosomes with a single TFBS were bound by a by a regulatory protein at increasing concentrations[15]. To examine different TFBSs or to determine their optimal translational or rotational settings within the nucleosome, it was necessary to repeat this assay for each TFBS and specific translational or rotational setting. An approach called NCAP-SELEX (<u>n</u>ucleosome <u>c</u>onsecutive <u>a</u>ffinity <u>p</u>urification–<u>s</u>ystematic <u>e</u>volution of ligands by <u>ex</u>ponential enrichment) was recently developed that uses randomized nucleosomal DNA libraries with TF-nucleosome enrichment followed by next-generation sequencing [16]. However, NCAP-SELEX is limited to short nucleosome fragments and cannot be used to directly compare binding to different locations in the nucleosome or to linker DNA. To compare TFBSs at various nucleosomal positions, competitive nucleosome-binding assays and Senu-seq combine electrophoretic mobility shift assays (EMSAs) with a nucleosome library and next-generation sequencing [4, 5, 17], but these approaches are limited in the number of sequences that can be examined and are restricted to experimentally defined nucleosome-positioning sequences (NPSs). To comprehensively examine TF-nucleosome binding across various NPSs and TFBSs and in *in vivo-*targeted nucleosomes (ITNs), we developed Pioneer-seq (transcri<u>p</u>tion fact<u>o</u>r <u>n</u>ucl<u>e</u>osom<u>e</u> binding protocol), which can be used to examine 7,500 nucleosomes in a single assay.

## Results

### Pioneer-seq workflow

Pioneer-seq expands on preexisting competitive nucleosome-binding assays by examining thousands of nucleosomes, which include nucleosomes based on highly characterized NPSs and nucleosomes based on ITNs. Pioneer-seq uses 5S rDNA [18], MMTV LTR [19], and the artificial synthetic sequence Widom 601 DNA as NPSs [20]. Use of these highly characterized NPSs is advantageous due to the availability of their defined structures and dynamics [15]. To analyze each TF that will be studied, one of its TFBSs is incorporated into each of the three NPSs at intervals of 1 base pair. This includes sites both within the core 147-bp nucleosome and outside in the linker regions. For each TF that will be studied, multiple TFBSs can be examined within a single experiment. Each NPS with the inserted TFBS is flanked by PCR primers (19–20 bp) for a total sequence length of 230 bp.

In addition to highly characterized NPSs, Pioneer-seq can also include genomic locations targeted *in vivo* by a TF of interest. These ITNs are defined by examining nucleosome positioning data from MNase-seq or NOMe-seq with TF binding data from ChIP-seq, ChIP-nexus, or Cut & Tag. The 191-bp sequences centered at the ITN are then filtered for predicted nucleosome formation ability [21] and the presence of the specific TFBS. In total, 7,500 sequences are designed for each library such that most sequences are not specific for any particular TF. All DNA sequences are synthesized as an Agilent 230-bp oligonucleotide library. Nucleosomes are then assembled with salt gradient dialysis using all nucleosome sequences simultaneously; free DNA is removed from the nucleotide library by a sucrose gradient.

The purified nucleosome library is used in binding assays in which the TF of interest is added at increasing concentrations. TF-nucleosome complexes are detected after a short incubation by separating the reaction mixtures on a native polyacrylamide gel (the first lane of which contains only nucleosomes to measure the background and input levels for each experimental replicate). Nucleosomes that are bound by the TF are identified by sequencing the DNA that is extracted and purified from shifted bands in the gel. The sequencing results are then analyzed and mapped back to the original 7,500-nucleosome library. Figure 1 illustrates the Pioneer-seq workflow.

**Fig 1.**
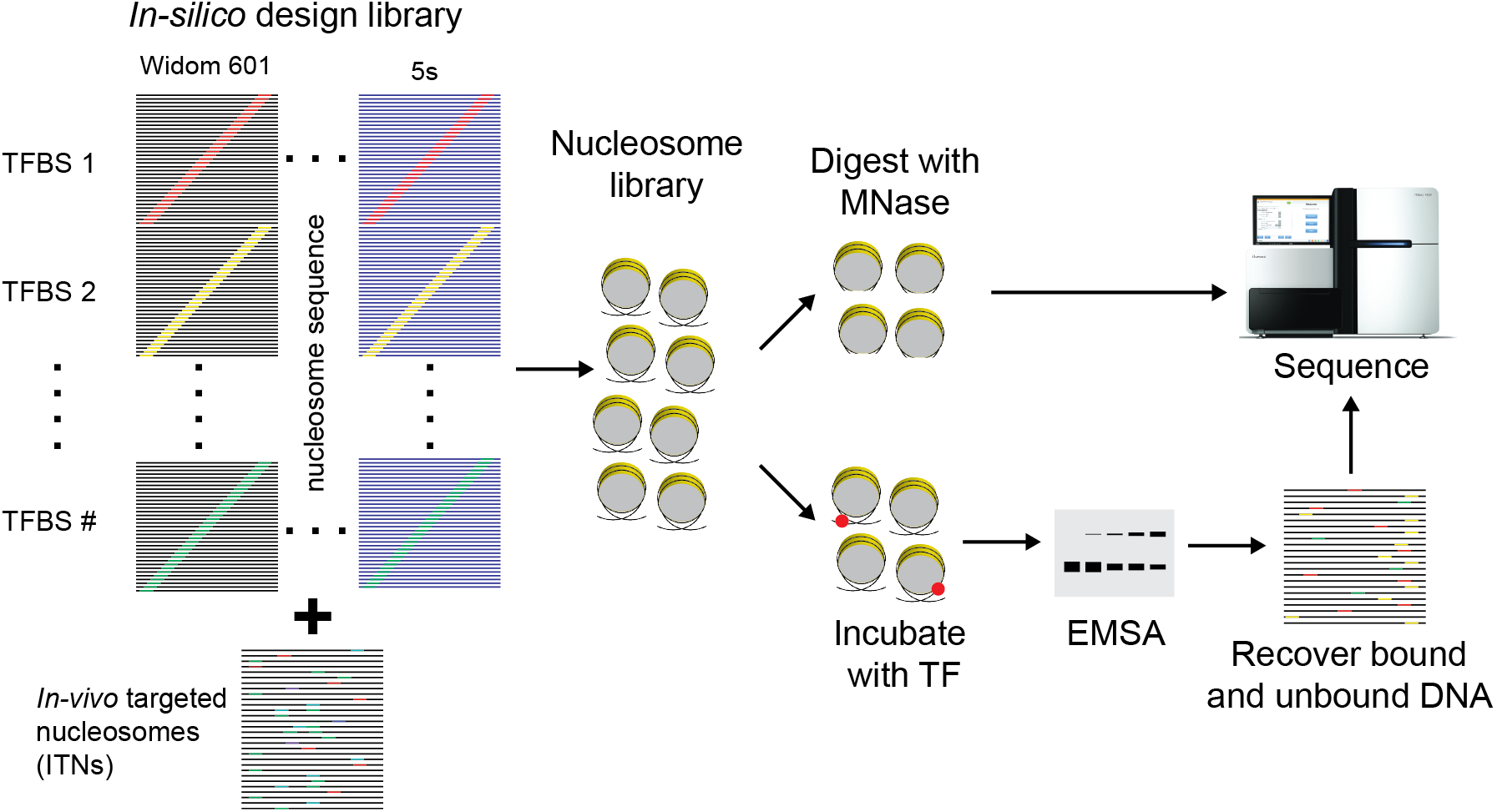
Overview of Pioneer-seq. With this method, 230-bp DNA sequences of interest are designed in batches of 7,500 sequences, including well-characterized nucleosome-positioning sequences (NPSs; Widom 601 or 5S rDNA) and sequences for *in vivo-*targeted nucleosomes (ITNs) (left). The method enables testing of binding-site variation and positioning, flanking-sequence content, and combinatorial binding events. Nucleosomes are formed and purified on all 7,500 sequences to generate a nucleosome library (middle). The entire nucleosome library is incubated with increasing amounts of a transcription factor (TF) of interest (bottom right). The TF-nucleosome complexes are separated by electrophoretic mobility shift assay (EMSA), and the bound and unbound DNA are recovered, quantified, and sequenced. Nucleosome positioning and accessibility for every DNA sequence in the nucleosome library are determined by digestion with micrococcal nuclease (MNase) (top right). The resulting DNA fragments are sequenced and mapped back to the initial library. TFBS, TF binding site.

TF-nucleosome binding is then determined by calculating the “relative supershift,” which compares a nucleosome sequence to a collection of non-specific sequences in the same binding experiment [5]. By comparing each nucleosome sequence to the non-specific control sequences from the same lane, loading, PCR amplification, next-generation sequencing, and binding to additional unknown motifs are all internally controlled.

Nucleosome formation and positioning for the Pioneer-seq library is then determined by MNase-seq [22, 23] . For these experiments, MNase-digested DNA from the nucleosome library is sequenced and mapped back to the 7,500-nucleosome library to determine which regions are protected; results for the well-defined NPSs included in this Pioneer-seq library are consistent with known nucleosome structures (Fig. S1).

### Defining TF-nucleosome binding

To evaluate Pioneer-seq, we examined nucleosome binding for the Yamanaka factors OCT4, SOX2, KLF4, and c-MYC [24]. OCT4, SOX2, and KLF4 have been shown to act as pioneer factors that directly bind to chromatin regions inaccessible to other TFs and subsequently trigger transcriptional competency by directing chromatin remodeling [25, 26]. Despite their pioneering capabilities, OCT4, SOX2, and KLF4 cannot bind to all their TFBS within a genome [27].

Pioneer-seq was first performed across a wide range of TF concentrations to determine the optimal signal for the assay (Fig. S2-5). The strongest binding occurs for all factors at TFBSs located outside of the nucleosome region within the linkers (Fig. S6). Naked libraries were also used for binding assays with the same conditions as performed for Pioneer-seq (Fig. S7). For all TFs tested in these naked-library binding assays, there was no predominant shifted gel band, with limited differences in binding between sequences. Within nucleosomes there is extensive binding variability depending on the NPS and TFBS. The Widom 601 NPS is the most extensively studied NPS and has been a model for studying nucleosome structures and dynamics [20]. In our experiments 601 nucleosomes are the most efficiently formed nucleosomes (Fig. S8). The binding of OCT4, SOX2, KLF4, and MYC to the 601 nucleosomes was inhibited when their TFBSs were near the nucleosome dyad (Fig. 2), similarly to that observed for TP53 and TP63 in previous studies [4, 5]. For KLF4 there were two TFBSs (with corresponding reverse complement sequences): Klf4-1 (CCCCACCC) is derived from the motif MA0039.4 from Jaspar [28], and Klf4-2 (GCCCCGCCCCGCCCC) is derived from the KLF4 long motif discovered in mouse embryonic stem cells [29]. KLF4 bound strongly to Klf4-1 only when the site was positioned >55 bp from the 601 dyad. Results for binding to Klf4-2 were noisier at some internal nucleosome positions because Klf4-2 appears to disrupt nucleosome formation. The results for the reverse complement sequences (Klf4-1RC and Klf4-2RC) are similar to those for the direct sequences. For OCT4, there is an OCT4 TFBS (TATGCAAAT) and a joint Oct4-Sox2 TFBS (CTTTGTTATGCAAAT). Binding of OCT4 to both TFBS is very similar, as the OCT4 target sequence is the same. However, there were small differences in their relative supershifts that resulted from differences in the center positioning of the TFBSs. For SOX2, there is a joint Oct4-Sox2 TFBS and a SOX2 TFBS (ACAATGG). SOX2 binds both sequences in the linkers and can bind the Oct4-Sox2 TFBS at the nucleosome edge. The non-pioneer factor MYC was examined with a single palindromic TFBS (ACCACGTGGT) derived from the motif MA0059.1 from JASPAR [28]. MYC can bind its TFBS in the linker and when its TFBS is located > 55bp from the dyad.

**Fig 2.**
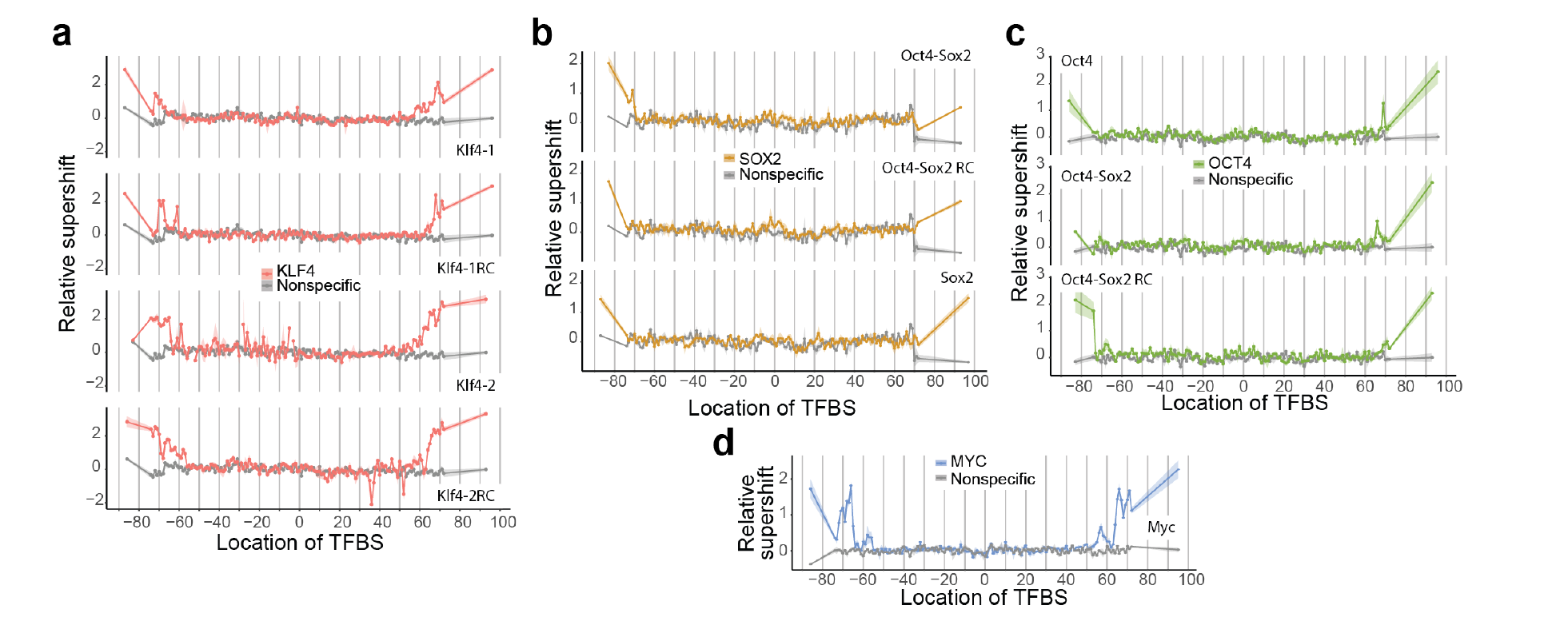
TF binding to the Widom 601 nucleosome. The specific and nonspecific TFBS is positioned across all possible locations along the 601 nucleosome with TFBSs in the left and right linkers to generate a total of 149 unique nucleosomes per TFBS. The relative supershift for each nucleosome is determined by counting the frequency of each sequence within the shifted band in the electrophoretic mobility shift assay and comparing it to that for nonspecific binding (i.e., binding to a sequence without the TFBS for that particular TF). This value is then normalized to the input ratio of nucleosomes (see Eq. 1). Shading around each line is SEM. Binding for KLF4 (**a**), SOX2 (**b**), OCT4 (**c**), and MYC (**d**) is shown for nucleosomes with their specific TFBSs along with binding to a nonspecific TFBS nucleosome sequence (shown in gray). Breaks in the trace for Klf4-2 TFBS indicate missing data as a result of inefficient nucleosome formation. RC, reverse complement.

With Pioneer-seq, we can compare the binding of TFs across different NPS in a single assay. We compared binding to nucleosomes based on the NPSs of Widom 601, 5S rDNA, and the MMTV long terminal repeat. KLF4, OCT4, SOX2, and MYC were able to bind to their TFBSs closer to the dyad for 5S nucleosomes than for 601 nucleosomes, with binding inhibited ±30 bp from the dyad. In addition, binding was asymmetric, with increased binding on the left side of the 5S nucleosome. Binding to the MMTV nucleosome was similar to that for the 5S nucleosome, with limited binding within 30 bp of the dyad. However, asymmetric binding occurred on the right face of the MMTV nucleosome. All TFs tested had greater binding to TFBSs in 5S and MMTV nucleosomes than to those in the 601 nucleosomes.

**Fig 3.**
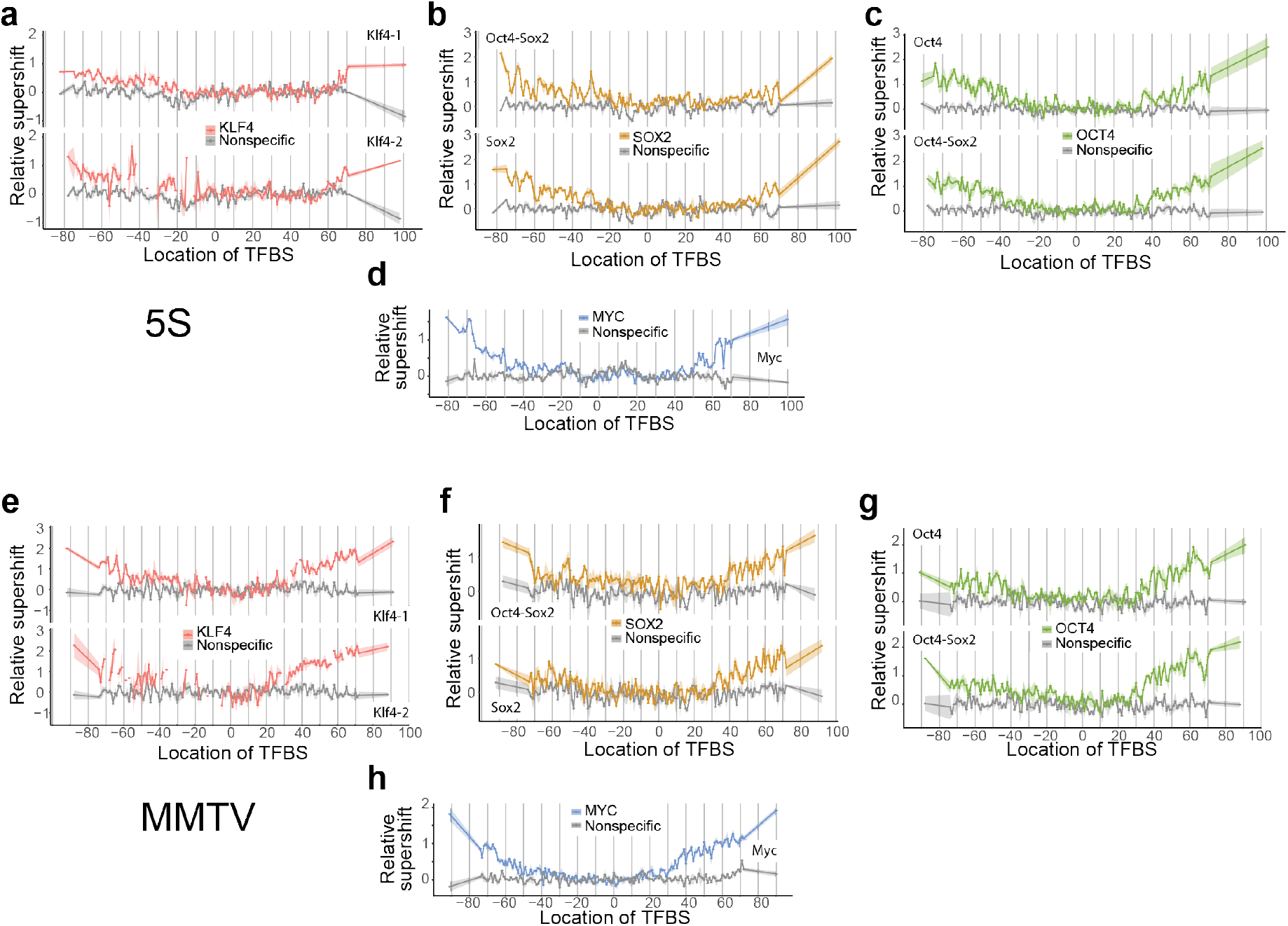
TF binding to 5S and MMTV nucleosomes. The TFBSs were positioned across all possible locations along the 5S (**a–d**) and MMTV (**e–h**) nucleosomes, with TFBSs in the left and right linkers. Relative supershifts for KLF4 (**a, e**), SOX2 (**b, f**), OCT4 (**c, g**), and MYC (**d, h**) are shown for their specific TFBSs along with binding to a nonspecific TFBS nucleosome sequence (shown in gray). Breaks in the trace for Klf4-2 TFBS in panels a and e indicate missing data as a result of inefficient nucleosome formation.

### Rotational and translational settings of nucleosome sequences drive binding differences

Because all 7,500 nucleosome sequences are exposed simultaneously to TFs in a single experiment, the binding to different NPSs or TFBSs can be directly compared. In general, we found that binding to TFBSs within the 601 nucleosome was strongly inhibited, whereas binding to TFBSs within the MMTV and 5S nucleosomes was less inhibited, and increased as TFBSs approached the edges of the MMTV and 5S nucleosomes. As shown in Fig. 4a, SOX2 binding to the OCT4-SOX2 TFBS differed depending on the NPS. SOX2 bound strongly to the left face of the 5S nucleosome, with periodic binding at -30 and -40 sites. SOX2 has a high-mobility group (HMG) box domain, which binds in the minor groove of DNA and induces helical bending [30]. The pre-bent conformation of nucleosomal DNA may thus facilitate the binding of SOX2 and other HMG domain-containing proteins [31].

**Fig. 4.**
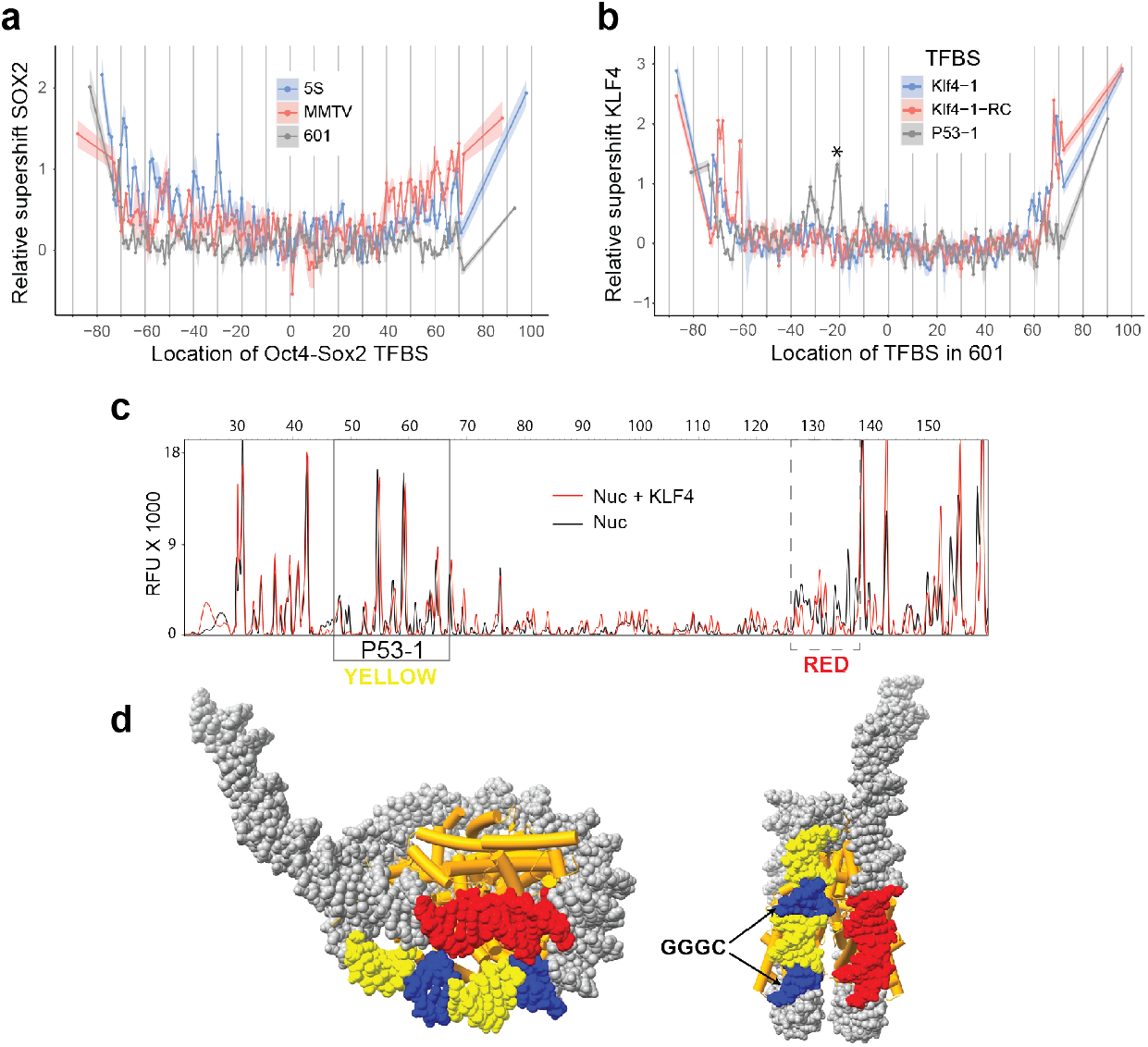
Nucleosome binding depends on nucleosome positioning sequence and binding site. **a**, Relative supershifts for SOX2 binding to the Oct4-Sox2 TFBS (CTTTGTTATGCAAAT) within the 5S, MMTV, and 601 nucleosomes. **B,** Relative supershifts for KLF4 binding to Klf4-1 (CCCCACCC), Klf4-1RC (GGGTGGGG), and p53-1 (GGGCATGTCCGGGCATGTCC) within the 601 nucleosome; “*” indicates the nucleosome validated in DNase-I footprinting experiments. RC, reverse complement. **c,** DNase-I footprinting of nucleosomes containing the p53-1 TFBS at SHL -2; an additional footprint observed on the neighboring gyre is indicated with the dashed-line box. Nuc, DNase-I digestion. **d,** Model of the 601 nucleosome with the p53-1 motif highlighted in yellow, the two GGGC in blue, and the additional footprinted region in red.

In addition to the KLF4 TFBSs in the library, KLF4 can bind to the TFBSs for p53. At these sequences, KLF4 can bind at the linkers and at specific rotational and translational settings near the nucleosome dyad. Its preferred TFBS is unbound when positioned at the same internal locations. The p53-1 and p53-2 TFBS are both bound by KLF4 in the linker regions but not as strong as the KLF4 TFBS (Fig. S6). On the other hand, when positioned near Super-Helix Location (SHL) -2 (-20 bp from dyad) within 601 both of the p53 motifs are bound (Fig 4b, S9). To confirm KLF4 binding to the p53-1 (GGGCATGTCCGGGCATGTCC) site on 601, we performed validation assays with DNase-I footprinting. In this experiment, the single nucleosome containing the p53-1 TFBS at SHL-2 is treated with DNase I before and after KLF4 binding (Fig 4c). Changes in protection are distinguishable at the p53-1 TFBS. KLF4 appears to be binding at two GGGC sequences spaced 10bp apart in the p53-1 TFBS. The two GGGC sequences are partially exposed in neighboring major grooves (Fig 4d). The DNase-I footprinting also showed an additional footprint located at SHL 5.5. This region shown in red in Fig 5d is directly next to the p53-1 TFBS located on the other DNA gyre, suggesting that KLF4 binding to p53-1 at SHL-2 disrupts the region near the entry/exit on the opposite side.

**Fig 5.**
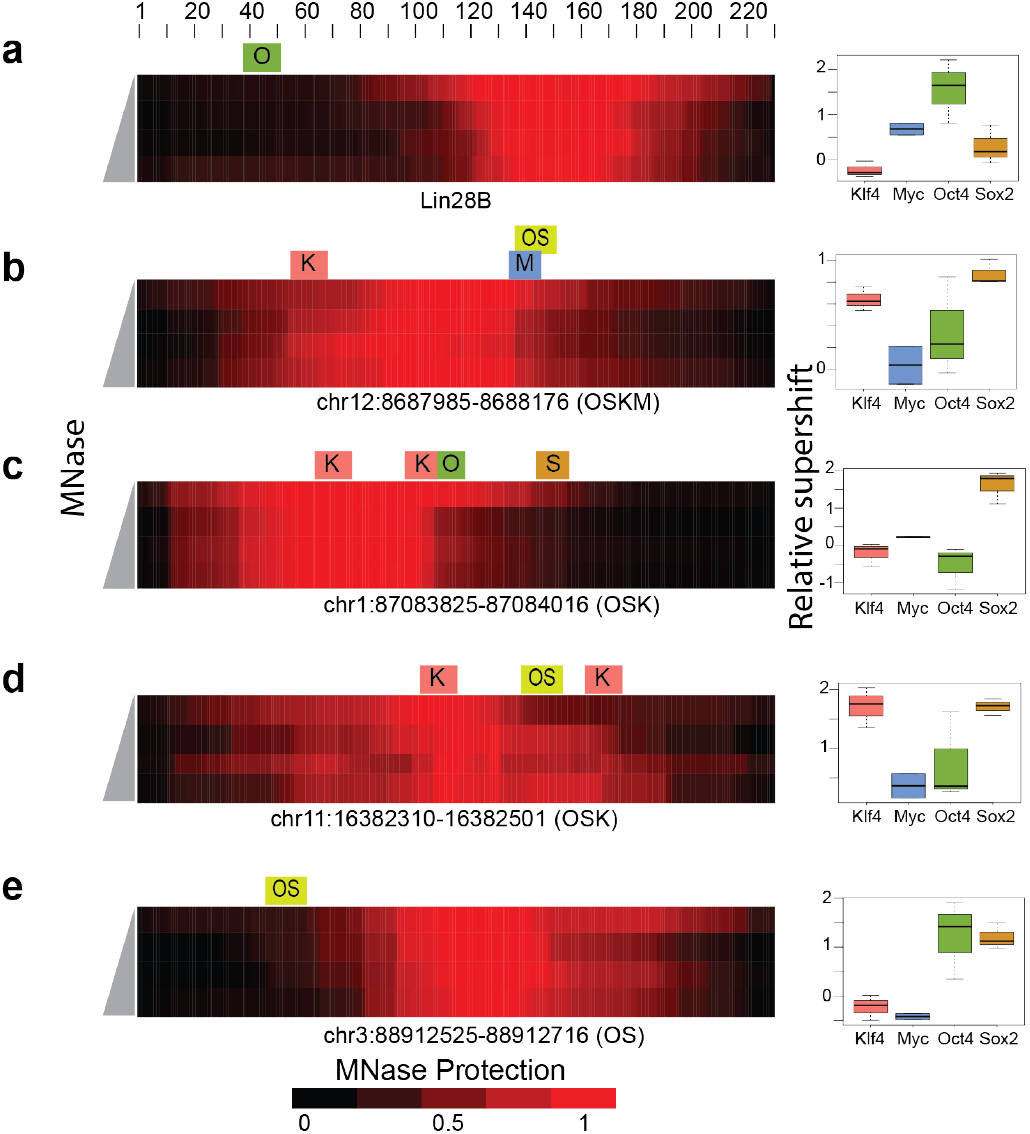
Binding to *in vivo*-targeted nucleosomes. The locations of TFBSs with MNase protection for *in vivo*-targeted nucleosomes (ITNs) are shown (red color scale at bottom). MNase protection was measured as the percentage of nucleosome bases that were protected from MNase digestion and calculated for each base pair as the ratio of base pair coverage to the total reads for that specific nucleosome: **a**, Lin28B nucleosome sequence bound with O; **b**, ITN bound with O, S, K, and M. **c–d**, ITN bound with O, S, and K; **e,** ITN bound with O and S. O, OCT4; S, SOX2; K, KLF4; M, MYC. The relative supershifts for each nucleosome are shown for KLF4, MYC, OCT4, and SOX2 binding on the right.

### Examining ITNs

In addition to well-characterized NPSs, ITNs at gene regulatory regions were included in the nucleosome library. These nucleosomes were identified using ChIP-seq binding data from induction of OCT4, SOX2, KLF4, and MYC in fibroblasts during IPSC generation [27]. Nucleosome positions were defined by NOMe-seq from the fibroblast cell line (IMR90) [32]. After filtering nucleosomes by predicted nucleosome formation efficiency, 372 nucleosomes were designed. In total, ∼20% of the ITNs failed to make stable nucleosomes for our assay. The remaining ITNs had a formation efficiency similar to 5S and MMTV nucleosomes (Fig. S8).

The Lin28B nucleosome is a ITN that has been examined by multiple groups [14]. We defined the nucleosome-protected region for Lin28B with MNase-seq and show a protected region centered at 150 bp with an OCT4 TFBS at position 38-46 bp. Pioneer-seq results show that only OCT4 can bind specifically to the Lin28B nucleosome. These results are consistent with DNase-footprinting and binding-specificity assays for KLF4, MYC, OCT4, and SOX2 [14]. The previously characterized ITNs located at ALBN1, NRCAM, CX3CR1, and ESRRB were also included within this library (Fig. S10). Other ITNs that were bound by OCT4, SOX2, KLF4, and MYC are shown in Fig. 5b–e.

By examining the Oct4-Sox2 sites, we can directly compare OCT4 and SOX2 binding at their TFBS when they are at the same nucleosome positions. At most Oct4-Sox2 TFBSs, SOX2 can bind, while OCT4 only binds when the TFBS is outside the protected nucleosomal region (Fig 5 b,d,e; S11). To compare binding between factors at ITNs, we have plotted the relative supershift for all ITNs containing a binding site. KLF4 and SOX2 bind to a larger percentage of their ITNs than MYC and OCT4 (Fig. 6a).

**Fig 6.**
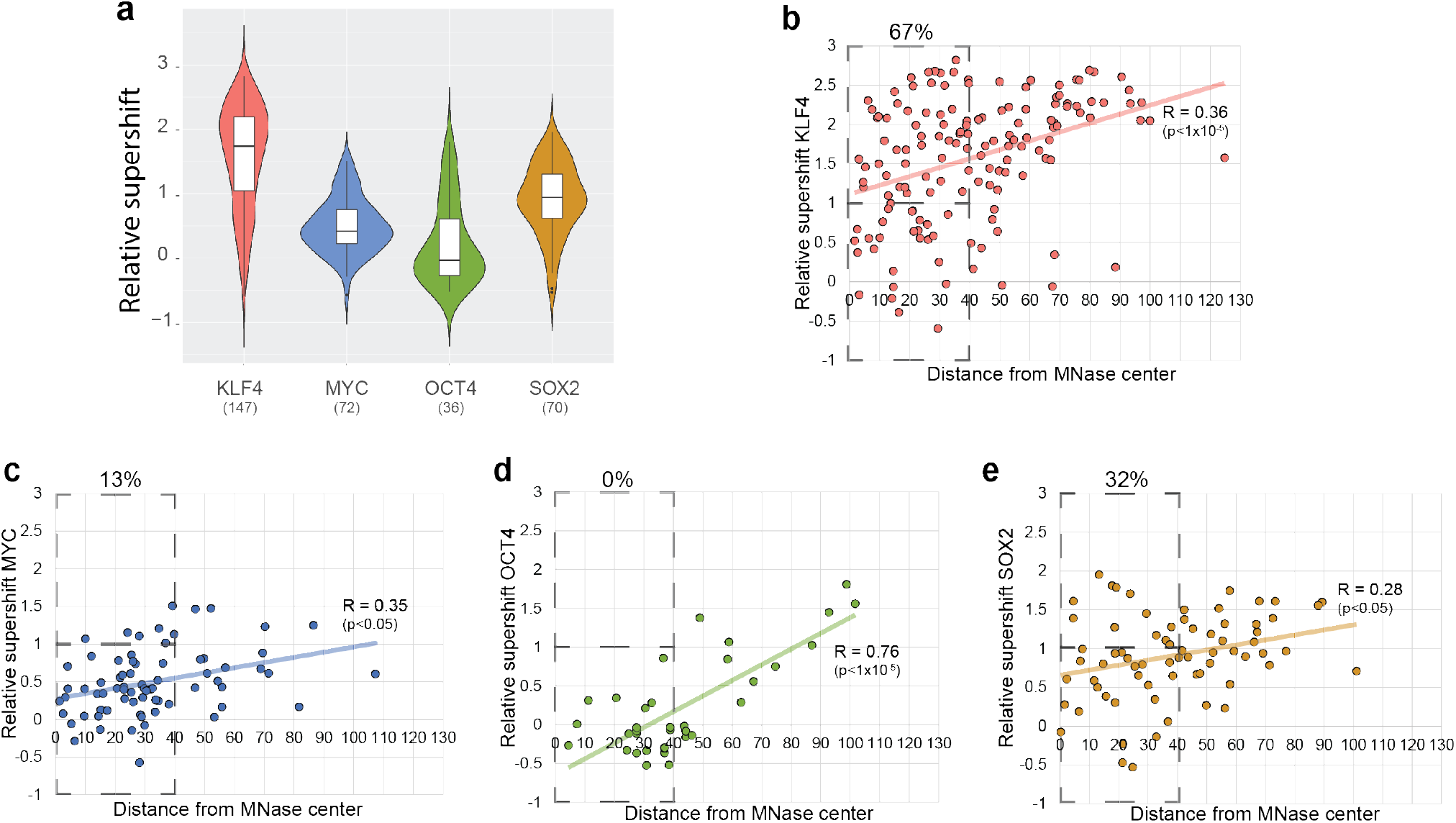
Binding to *in vivo-*targeted nucleosomes. **a**, Violin plot showing relative supershifts for KLF4, MYC, OCT4, and SOX2 binding to *in vivo-*targeted nucleosomes (ITNs) containing their transcription factor binding sites (TFBSs). **b–e,** Relative supershifts for binding to ITNs compared to the distance of the TFBS distance from the center of MNase protection.

To understand the role of nucleosome positioning for the ITNs, we mapped the TFBS in relationship to the center of MNase protection (Fig. 6b-e). Binding of KLF4, OCT4, SOX2, and MYC are significantly correlated with the distance of their TFBSs from MNase-protection centers. These results are very similar to the results for 5S and MMTV, suggesting that distance to dyad is a major factor regulating TF-nucleosome binding. To examine binding near the nucleosome center we selected the region 40 bp from the MNase center and determined the percentage of bound sites to the total number of sites within 40 bp. KLF4 bound 67%, SOX2 32%, MYC 13%, and OCT4 0%. This suggests that KLF4 and SOX2 have a special ability to target some of their TFBSs within a nucleosome and support the results with NPSs showing that KLF4 bound at SHL -2 and -3 and SOX2 bound at SHL -3 and -4.

## DISCUSSION

Pioneer factors are a proposed class of TFs that can bind inaccessible genomic regions and then facilitate the binding of other TFs. This ability is attributed to specific DNA-binding domains that are able to target their binding sites within a nucleosome [31] and bind to partial motifs that are accessible on the nucleosome surface [14]. However, recent studies with the archetypal pioneer factor FOXA1 and its non-pioneer cofactor HNF4A have shown that binding is defined by the TFBS, specifically its density and affinity, rather than differences in TFs themselves[33]. Our results suggest a more complex model.

The TFs we tested bound most strongly at the nucleosome edge, but this differed according to the NPS being bound. These results are consistent with a dynamic partial unwrapping of DNA from histones at sites where the DNA enters or exits the nucleosome, exposing the DNA to TFs [34, 35]. Edge binding would be heavily dependent on TFBS affinity and the number of sites. We also found that two of the TFs (KLF4 and SOX2) targeted sites close to the nucleosome dyad at specific rotational settings. By using ITNs, we were able to show that KLF4 and SOX2 behave differently than OCT4 and can target their binding sites near the nucleosome center.

KLF4 appears to be binding to two partial motifs (CCCG) located in consecutive partially exposed major grooves, similar to the nucleosome binding of another zinc-finger TF, GATA3. GATA3 binds to split 5′-GAT-3′ motifs in solvent-exposed major grooves [36]. KLF4 will only bind the non-typical motif at these internal locations, which suggests that the partial motif or neighboring bases provide a structural binding context that is missing for other TFBSs. The binding of KLF4 to the internal p53 sites in the 601 NPS but not the other NPS suggests that nucleosome structure also facilitates certain binding events. Although the overall nucleosome structure is consistent across different NPS, certain patterns of dinucleotides cause differences in the groove width and helical deformation of nucleosomal DNA [37, 38]. Indeed, groove width and the extent of helical deformation vary substantially among different sites with the same rotational settings [39, 40]. Single-molecule DNA-unzipping experiments have shown that position-specific histone-DNA interactions also vary across the nucleosome [41]. The binding of KLF4 to p53 TFBS may have biological significance. KLF4 with the p53-family member p63 coordinate gene regulation during skin development and have been shown to jointly target super enhancers [42].

There are still many questions about how TFs can bind nucleosomal DNA that can be addressed with Pioneer-seq. The vast majority of studies on nucleosomes have used the Widom-601 NPS. This sequence is extremely well-studied and is the sequence used for 114 structural studies. Widom 601 forms nucleosomes *in vitro* very efficiently in a single predominate position allowing reproducible and well-defined structures [15]. 5S and MMTV nucleosomes are biologically derived sequences that have been used for various studies [37, 43]. 5S and MMTV nucleosomes are not as stable as 601 and form with efficiencies similar to ITNs (Fig. S8). In this study, the nucleosome-binding abilities of each TF depended on the NPS being bound. In general, 601 nucleosomes were strongly inhibitory for TF binding, whereas 5S and MMTV nucleosomes were only inhibitory when the TFBSs were close to the nucleosome dyad. These results suggest that TF binding may be best understood by examining both ITNs and model nucleosomes. Use of ITNs does have limitations, because exact positioning of the dyad is unknown and weak nucleosome formation could limit specific sequences from being tested. For Pioneer-seq, we propose the use of ITNs along with a model nucleosome in which the TFBS can be positioned in all possible nucleosomal locations.

The ability for some TFs to bind a specific nucleosome site appears to depend on the binding sequence. This suggests that certain sites that require a pioneer factor may also have an alternative motif. The bases flanking the motif could impact nucleosome binding by affecting the structural presentation of the TFBS. TF binding could also be affected by events that influence the shape of the DNA around the nucleosome, such as DNA methylation, histone variants, and histone modifications. In the future, Pioneer-seq can be used to investigate these possibilities.

Lastly, transcription is regulated by a complex of multiple TFs that bind proximal regulatory regions. These multi-TF binding events can be directly cooperative, as seen when TFs physically interact, or indirectly cooperative to displace the nucleosome [44]. Pioneer-seq is ideally suited to test various models of cooperativity between TFs and enable the mechanistic dissection of these crucial regulatory events.

## Conclusion

In summary, Pioneer-seq is a powerful method for investigating the essential first step in gene regulation, the binding of TFs to inaccessible DNA located within nucleosomes. Due to its nature of comparing specific binding to non-specific binding across a whole nucleosome library, Pioneer-seq allows the direct comparison of sites located in various nucleosome positions, with differing NPSs, and with differing TFBSs. Pioneer-seq can be applied to directly address various mechanistic models for TF-nucleosome binding and can be used to uncover inherent TF-interaction differences.

## Methods

### Pioneer-seq library design

Three nucleosome positioning sequences (NPS), namely, Widom 601, 5S rDNA, and mouse mammary tumor virus (MMTV)-A, were used to form stable nucleosomes. Each sequence has been characterized by multiple biochemical assays [45] and was scanned for the presence of binding sites for the transcription factors (TFs) of interest using FIMO; sequences were modified to remove the binding sites [46]. Sequences were generated with the TF binding site (TFBS) of interest placed at every base pair positing in the nucleosome, with sites in both linker regions. In total, 149 sequences were designed for every TFBS with each NPS.

*In vivo*-targeted nucleosomes (ITNs) were determined by integrating datasets from chromatin immunoprecipitation with sequencing (ChIP-seq) and nucleosome positioning datasets. Locations bound by OCT4, SOX2, KLF4, or MYC were determined from ChIP-seq datasets [27]. Bound sites were checked for the specific TFBS, and locations lacking an identifiable TFBS were removed. Nucleosome positions were determined from NOMe-seq (nucleosome occupancy and methylome sequencing) from IMR90 cells (GSM543311) using the DANPOS algorithm [32, 47]. The position serving as the nucleosome center was then expanded to 191 bp and evaluated with a nucleosome scoring function [21]. The probability that the center base is part of a nucleosome was used as the probability score for each nucleosome. Nucleosomes with a score of <0.7 were removed from the design.

### Nucleosome library assembly

All nucleosome sequences were flanked by primer sequences to generate 230-bp sequences. The nucleosome library containing a total of 7,500 unique sequences was acquired from Agilent as a custom oligonucleotide library, which was amplified using Herculase II Fusion DNA polymerase in 100-µl reaction mixtures (1× Herculase II reaction buffer, 1 mM dNTPs, 200 pM Agilent library, 250 nM forward and reverse primers) with 15 PCR cycles. For a typical experiment, the DNA obtained from 11 reactions was purified with a QIAquick PCR purification kit (cat. no. 28104; Qiagen) and quantified with a NanoDrop spectrometer; fragment size was confirmed with a 2% agarose gel. Nucleosomes were then generated from H2A/H2B dimers and H3.1/H4 tetramers (NEB) by incubating the DNA sequences and the histones at an octamer/DNA molar ratio of 1:1.2 (in a solution containing 10 mM dithiothreitol [DTT] and 1.8 M NaCl) for 30 min at room temperature. The reaction mixture was transferred to a Slide-A-Lyzer MINI dialysis unit (10,000 MWCO, cat. no. 69750; Thermo Scientific). Dialysis was performed with 1.2 ml of the dialysis buffers at 4°C in 1.0 M NaCl for 2 h, 0.8 M NaCl for 2 h, 0.6 NaCl for 2 h, and TE buffer (pH 8.0) overnight at 4°C. Nucleosomes were then transferred to a clean 1.5-ml tube pretreated with 0.3 mg/ml bovine serum albumin (BSA). Nucleosome formation was then confirmed by 4% native polyacrylamide gel electrophoresis. Free DNA was removed from nucleosomes by using a 7%–20% sucrose gradient, and nucleosomes were concentrated and quantified via qPCR [4, 5]. Nucleosomes were then stored at 4 °C for up to 1 month.

### DNA binding assay followed by EMSA

The protein-nucleosome binding assays were carried out by incubating the purified nucleosome libraries described above and human full-length KLF4 (Origene TP306691), OCT4 (Origene TP311998), SOX2 (Origene TP300757), and MYC (Origene TP307611) with MAX (Origene TP306812) (in 7 µl DNA binding buffer (10 mM Tris-Cl [pH7.5], 50 mM NaCl, 1 mM DTT, 0.25 mg/ml BSA, 2 mM MgCl_2_, 0.025% Nonidet P-40, and 5% glycerol) for 10 min on ice and then 30 min at room temperature. Increasing concentrations of TF (0 to 3.2 pmol) were added to 0.2 pmol purified nucleosomes. Protein binding was detected by electrophoretic mobility shift assays (EMSAs) on 4% (w/v) native polyacrylamide gels (acrylamide/bisacrylamide, 29:1 [w/w], 7 × 10 cm) in 0.5× Tris-borate-EDTA buffer at 100 V at 4 °C. Initial EMSA experiments are done to across a wide range of TF concentrations to determine the optimal TF amount and to ensure the supershift is observed on the gel (Fig S2-5).

### DNA isolation and purification

After electrophoresis, DNA was imaged by staining with SYBR green (LONZA). All visual bands as well as the bands at the same locations in the other lanes were excised from the gel. The chopped gel slices were soaked in diffusion buffer (0.5 M ammonium acetate, 10 mM magnesium acetate, 1 mM EDTA [pH 8.0], 0.1% SDS) and incubated at 50 °C overnight. The supernatant was collected, residual polyacrylamide was removed with glass wool, and the DNA was purified with QIAquick spin columns (Qiagen). The DNA concentration for each sample was determined by qPCR using a standard curve generated from a control sequence.

### Library construction and sequencing

Illumina sequencing libraries were generated using a two-step PCR method, with 8 to 12 cycles of amplification for the first step including four sets of primers designed to offset sequence reads and dephase the libraries during Illumina sequencing (Extended Data Table 1). The number of cycles for the first-round PCR was determined using the sample concentration determined by qPCR. Each sample was then indexed using Nextera dual indices (Nextera XT index primer 1 [N7xx] and Nextera XT index primer 2 [S5xx]). After each PCR, reaction mixtures were cleaned up with AMPure XP beads (Beckman Coulter). The concentration of each sample was determined using the Invitrogen Quant-iT dsDNA assay kit, and equal amounts of each sample DNA were pooled and sequenced on an Illumina NextSeq 2×150. Sequencing and quality control were performed at the University at Buffalo Genomics and Bioinformatics Core.

### Pioneer-seq analysis

Illumina sequence reads were processed with an automated Snakemake pipeline of applications to refine and identify the sequences present in the sample pool [48]. The 3′ ends of Illumina FASTQ reads with low-quality scores were removed with Cutadapt using a quality cutoff of 30 (-q 30)[49]. Forward and reverse FASTQC reads were merged with Vsearch (--fastq_mergepairs) only if they shared at least 20 overlapping nucleotides (--fastq_minovlen 20) and had no more than two mismatched nucleotides between them (--fastq_maxdiffs 2)[50]. Primer sequences present at the ends of FASTQ reads were removed with Cutadapt. FASTQ reads of >220 nucleotides (nt) or <174 nt were filtered out with Cutadapt (--maximum-length 220 --minimum-length 174)[49]. FASTQ reads were converted to FASTA format using the FASTX-Toolkit FASTQ-to-FASTA command[51]. FASTA reads were mapped to a sequence in the reference library of 7,500 nucleosome sequences with Vsearch (--dbmatched) only if they had alignment lengths of at least 150 nt (--mincols 150), had at least 98.5% similarity (--id 0.985), and were the query and database sequence pairing with the highest percentage of identity (--top_hits_only)[50]. The results were then analyzed relative to control/nonspecific binding (relative supershift).

Relative supershift was determined from the supershift bands and controls for technical variability introduced by gel excision, PCR, library construction, or sequencing. In this method, each specific nucleosome sequence is measured relative to that for nonspecific binding (nucleosomes lacking a TFBS):

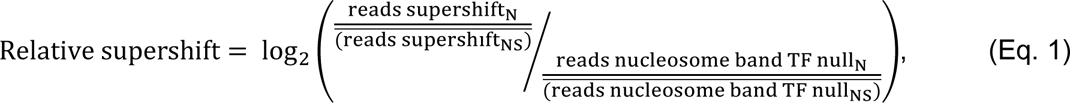

where N is one of the 7,500 nucleosome sequences, NS is the control nucleosome sequences, “reads supershift” is the supershift band, and “reads nucleosome band TF null” is the nucleosome band in the TF-null lane. The control non-specific nucleosome sequences are selected from the same NPS and are a collection of sequences that have different TFBSs. The non-specific TFBS are checked for binding to the particular TF in the linker regions. For each TF there are over 500 control sequences for 601, and 298 control sequences for 5S and MMTV.

Pioneer-seq was performed with multiple concentrations of TFs because of differences in inherent binding affinities and protein purity. An initial analysis of linker binding events was used to (i) confirm specific binding of the TF of interest and (ii) define the TF concentration with the largest binding signal compared to that for nonspecific binding. The relative supershift for a single TF concentration is presented throughout this manuscript: 0.4 pmol for KLF4, 0.2 pmol for OCT4 and SOX2, and 0.8 pmol for MYC/MAX.

### MNase-seq on nucleosome library

Nucleosome positioning for each sequencing in the library was determined with MNase-seq as previously described [52]. The nucleosome library (0.2 pmol/μl) was digested by MNase (0.05 U/μl) in nuclease digestion buffer (10 mM Tris-HCL [pH 8.0], 2 mM CaCl_2_) over a time course (0 to 25 min) at 37 °C; digestion was stopped with 2% SDS and 40 mM EDTA). Each sample was then incubated with proteinase K (16 μg) for 1 h at 55 °C. The DNA was purified from the reaction and concentrated with the QIAquick PCR purification kit. The concentrations of DNA in each sample were determined with the Invitrogen Quant-iT dsDNA assay kit and equalized. Illumina sequencing libraries were generated using an NEBNext Ultra II DNA library prep kit. Individual samples were multiplexed and sequenced via Illumina MiSeq 2×150.

MNase-seq results were quality filtered (*q* > 30) and adapter trimmed using Cutadapt [53]. The quality reads were merged and mapped to the 7,500 nucleosome library sequences using Vsearch[54]. The read counts and end positions were used to measure MNase protection, which was calculated for each base pair as the ratio of base pair coverage to total reads for that specific nucleosome.

### DNase-I footprinting

Nucleosomal DNA 5′ labeled with FAM (6-carboxyfluorescein) was formed into nucleosomes and purified as described above. Nucleosomes (50 ng) were bound with 0.4 pmol of KLF4 in DNA binding buffer (10 mM Tris-HCL [pH 7.5], 1 mM MgCl_2_, 10 µM ZnCl_2_, 1 mM DTT, 10 mM KCl, 0.5 mg BSA, 5% glycerol) at room temp for 1 h. Each sample was then incubated with 0.06 U DNase I in 50 µl of digestion buffer (10 mM MgCl_2_, 5 mM ZnCl_2_) at 25 °C for 1 min; digestion was stopped with 90 µl stop solution (NaCl_2_, 30 mM EDTA, 1% SDS). Digested DNA was then purified using phenol/chloroform/isoamyl alcohol and submitted to Roswell Park Genomic Facility for capillary electrophoresis fragment analysis on an ABI PRISM 3130xl Genetic Analyzer. The resulting data were analyzed using the Microsatellite analysis app from Thermo Fisher Scientific.

### Modeling of a KLF4-bound nucleosome

The structure for the Widom 601 nucleosome [55] was retrieved from the Protein Data Bank (PDB)[56] (PDB identifier 5OXV). The location of the KLF4-bound motif was determined from the Pioneer-seq results, and the structural residues were colored using ChimeraX software [57].

## Data availability

All Pioneer-seq results are stored on NCBI SRA at accession PRJNA892950.

