## Supplementary material for "Defining transcription factor nucleosome binding with Pioneer-seq": S1

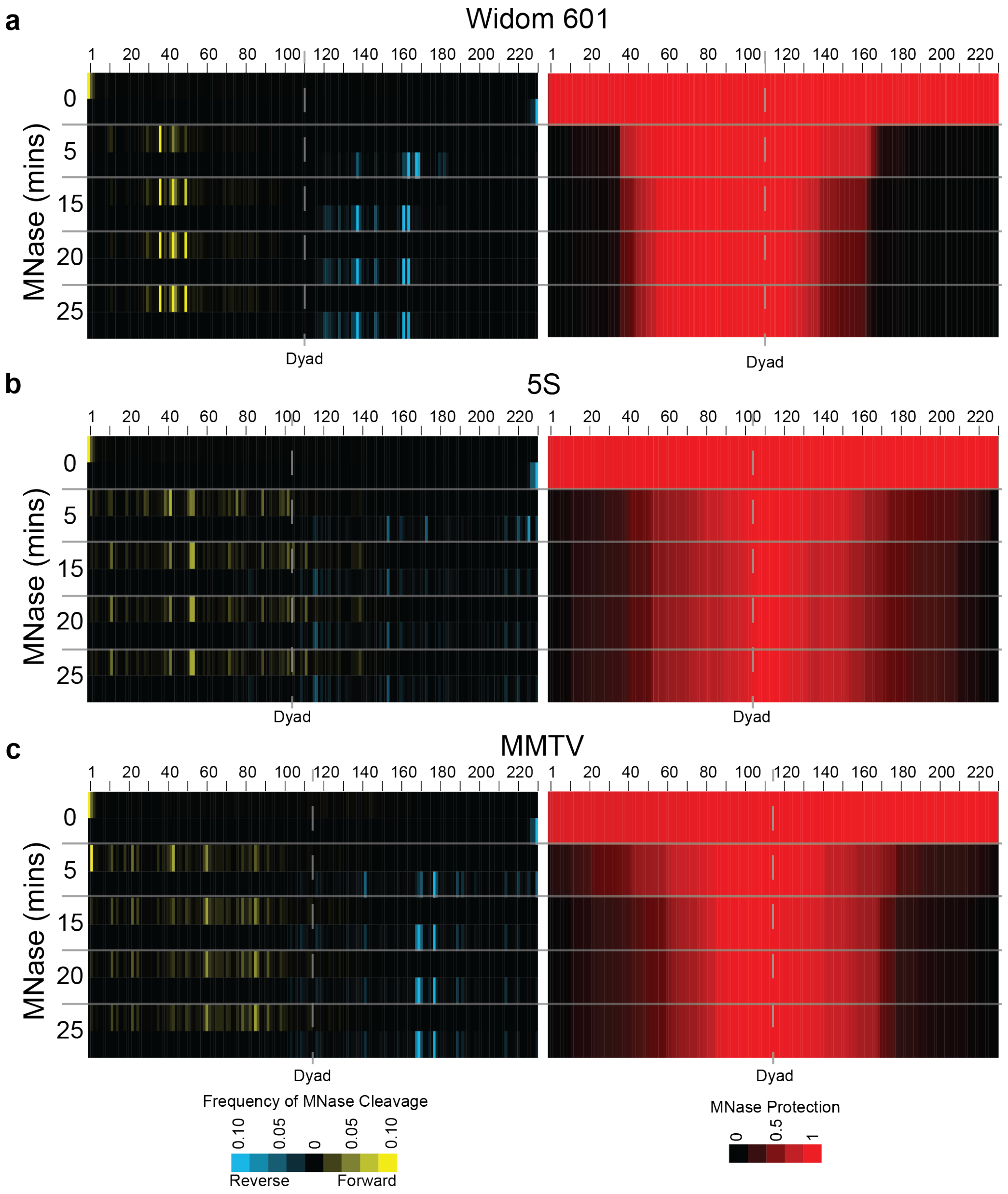


**Fig. S1. MNase-seq on nucleosome library.** Nucleosome libraries were digested with micrococcal nuclease (MNase) for various times. Sequence reads were then mapped back to a database of the 7500 sequences in the library. Then sequences from the same NPS (601, 5S, MMTV) were pooled together. Mapped fragment ends is used to determine frequency of MNase cleavage at specific bases (left). MNase protection is determined as the ratio of base pair coverage to the total number reads for that specific nucleosome (right). **(a)** Widom 601 nucleosomes, **(b)** 5S nucleosomes, **(c)** MMTV nucleosomes.

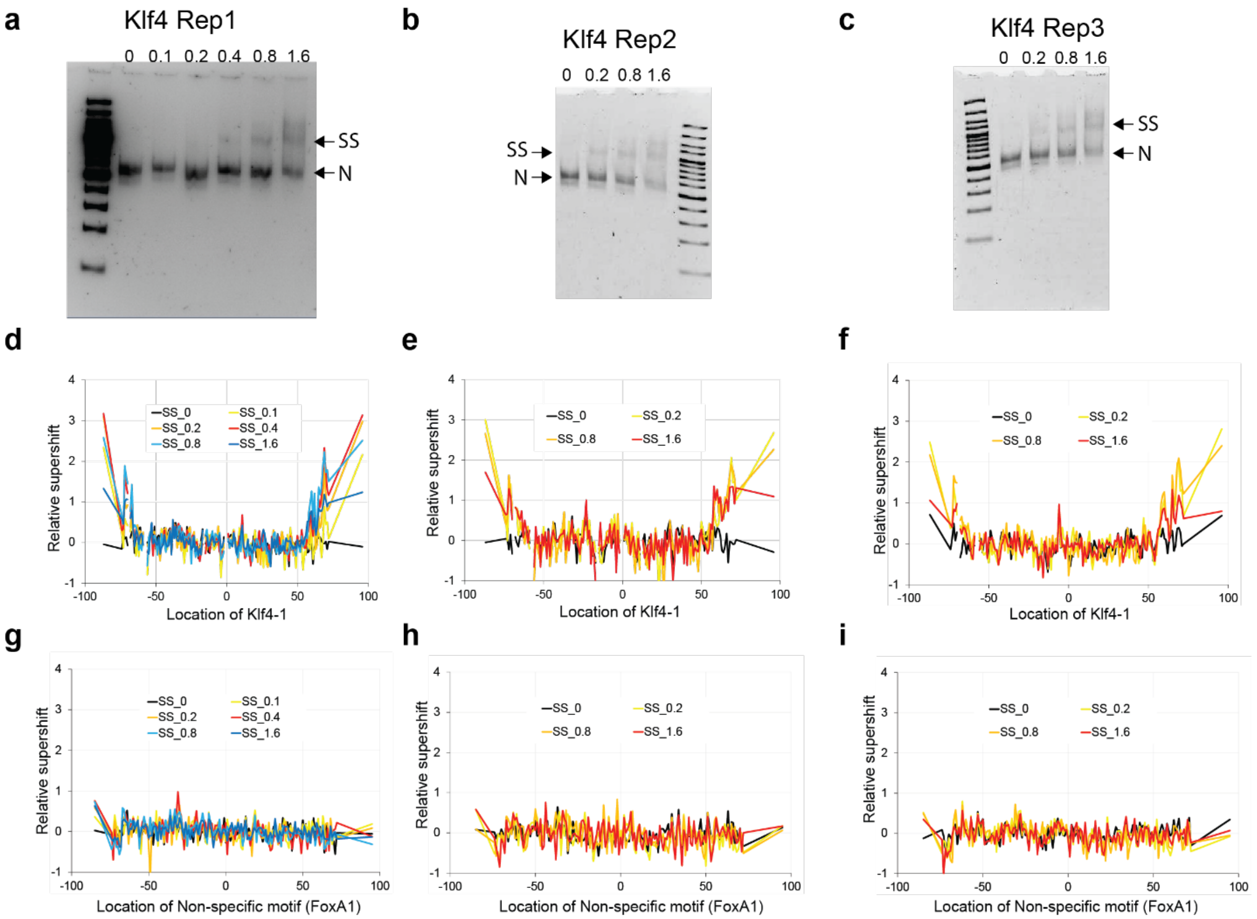


**Fig. S2. KLF4 Pioneer-seq binding assays.** (**a**,**b**,**c**) 7500 nucleosome sequences were bound to increasing amounts of KLF4 and separated by native PAGE. All assay lanes contain 0.2 pmol nucleosomes with 0, 0.1, 0.2, 0.4, 0.8 or 1.6 pmols of KLF4. Nucleosome and the supershift (SS) bands are indicated. (**d,e,f**) Relative supershift for KLF4 binding to the KLF4-1 TFBS (CCCCACCC) at all TF concentrations. (**g,h,i**) Relative supershift for KLF4 binding to the non-specific TFBS (TGTTTACTTTG) at all TF concentrations.

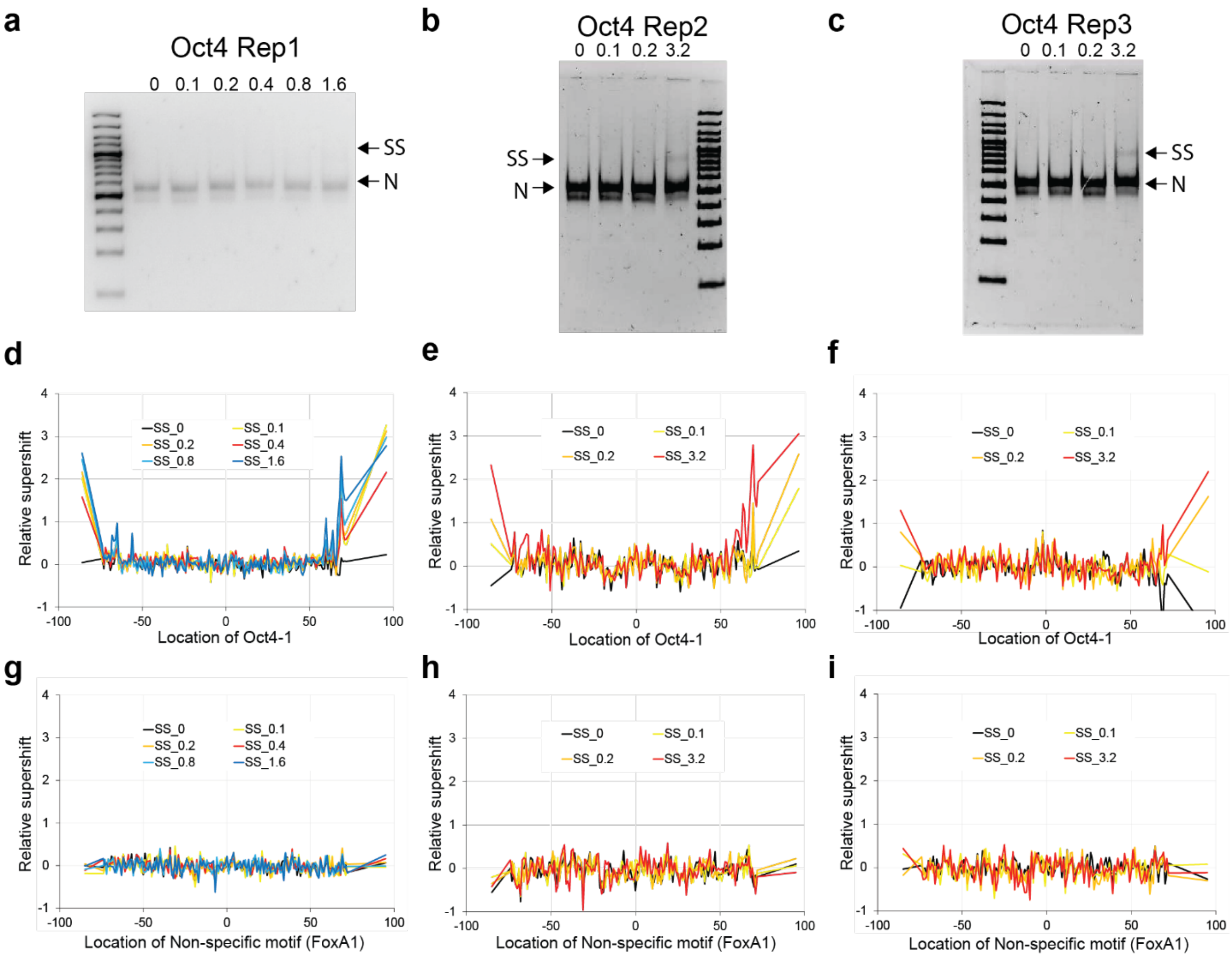


**Fig. S3. Oct4 Pioneer-seq binding assays.** (**a**,**b**,**c**) 7500 nucleosome sequences were bound to increasing amounts of OCT4 and separated by native PAGE. All assay lanes contain 0.2 pmol nucleosomes with 0, 0.1, 0.2, 0.4, 0.8, 1.6, or 3.2 pmols of OCT4. Nucleosome and the supershift (SS) bands are indicated. (**d**,**e**,**f**) Relative supershift for OCT4 binding to the OCT4-1 TFBS (TATGCAAAT) at all TF concentrations. (**g**,**h**,**i**) Relative supershift for OCT4 binding to the non-specific TFBS (TGTTTACTTTG) at all TF concentrations.

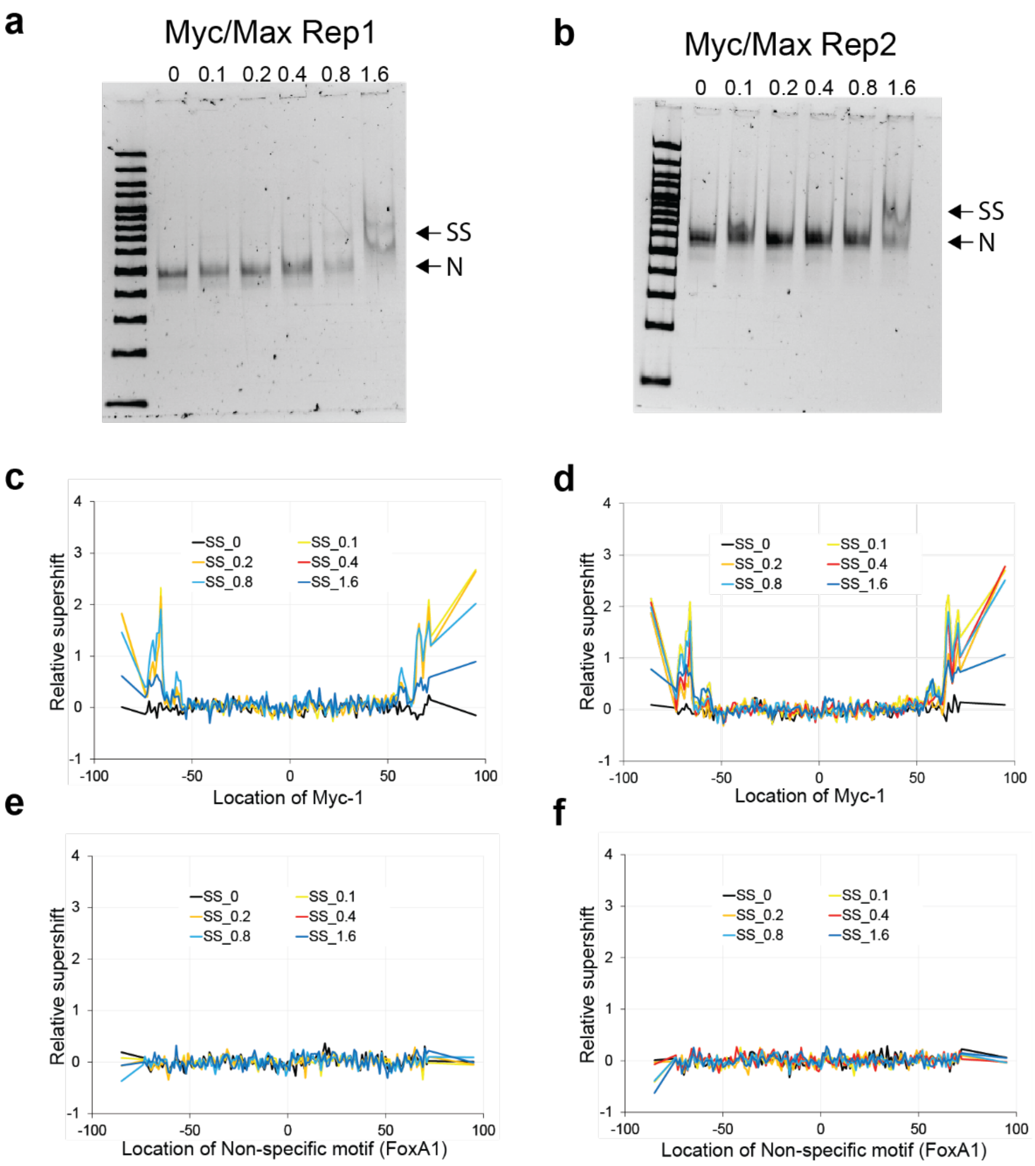


**Fig. S4. MYC/MAX Pioneer-seq binding assays.** (**a**,**b**) 7500 nucleosome sequences were bound to increasing amounts of OCT4 and separated by native PAGE. All assay lanes contain 0.2 pmol nucleosomes with 0.1, 0.2, 0.4, 0.8 or 1.6 pmols of MYC/MAX. Nucleosome and the supershift (SS) bands are indicated. (**c**,**d**) Relative supershift for MYC/MAX binding to the Myc-1 TFBS (ACCACGTGGT) at all TF concentrations. (**e**,**f**) Relative supershift for MYC/MAX binding to the non-specific TFBS (TGTTTACTTTG) at all TF concentrations.

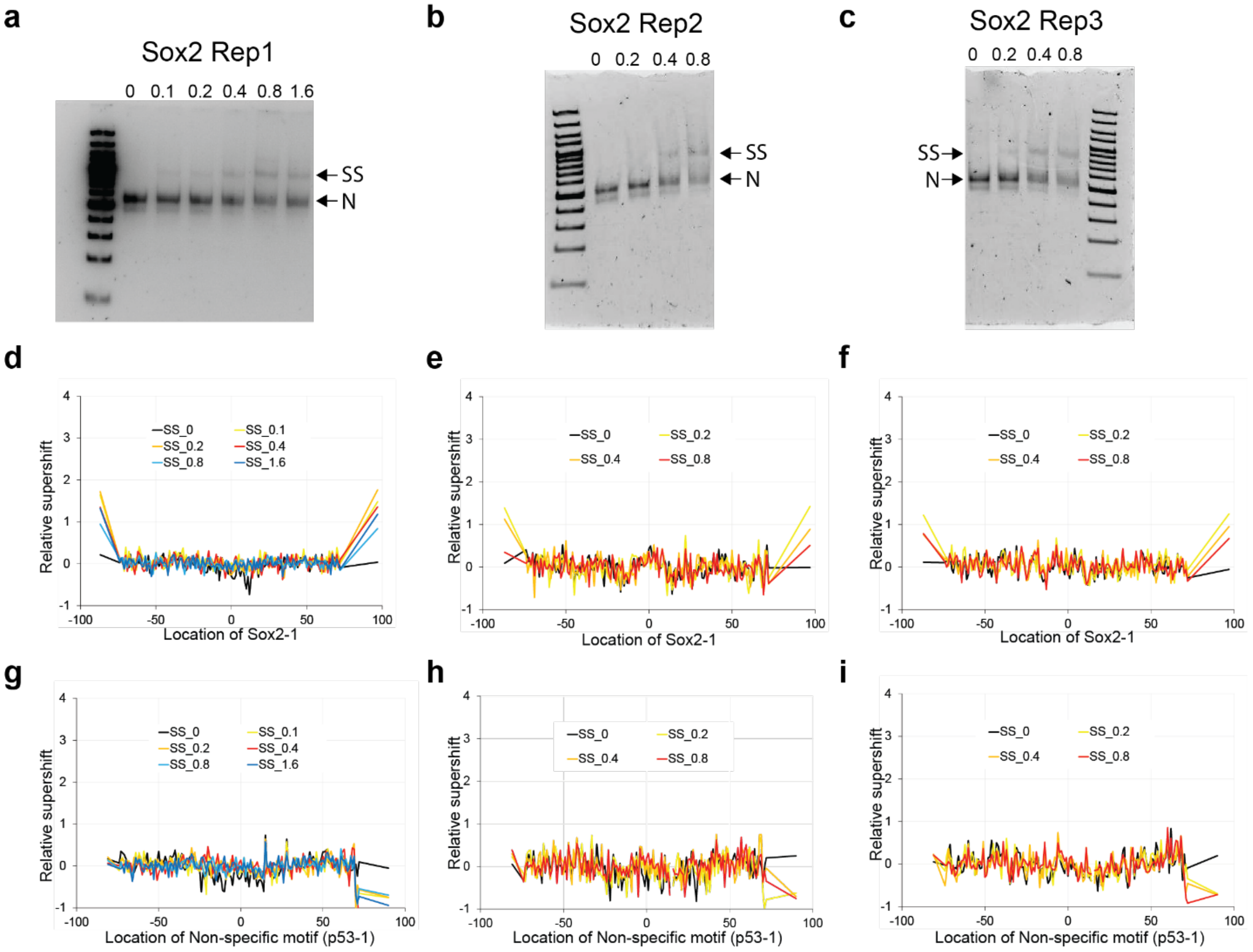


**Fig. S5. Sox2 Pioneer-seq binding assays.** (**a**,**b**,**c**) 7500 nucleosome sequences were bound to increasing amounts of SOX2 and separated by native PAGE. All assay lanes contain 0.2 pmol nucleosomes with 0, 0.1, 0.2, 0.4, 0.8 or 1.6 pmols of SOX2. Nucleosome and the supershift (SS) bands are indicated. (**d,e,f**) Relative supershift for SOX2 binding to the SOX2-1 TFBS (ACAATGG) at all TF concentrations. (**g,h,i**) Relative supershift for SOX2 binding to the non-specific TFBS (GGGCATGTCCGGGCATGTCC) at all TF concentrations.

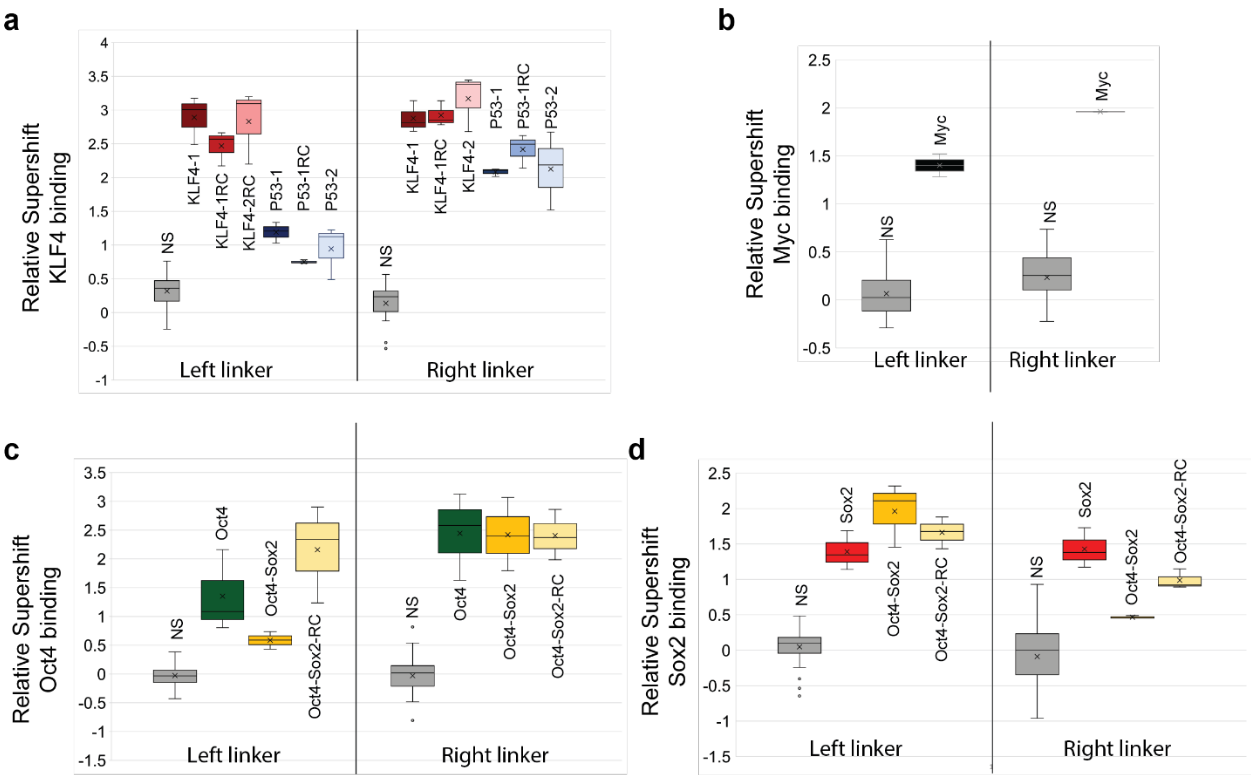


**Fig. S6. Binding at linker sites in 601 NPS.** Binding of **(a)** KLF4, **(b)** MYC, **(c)** OCT4, and **(d)** SOX2 to TFBSs located in the left and right linkers of the Widom-601 NPS (that is, TFBSs located outside the 147-bp nucleosome core). For every experiment a non-specific (NS) TFBS is shown for comparison.

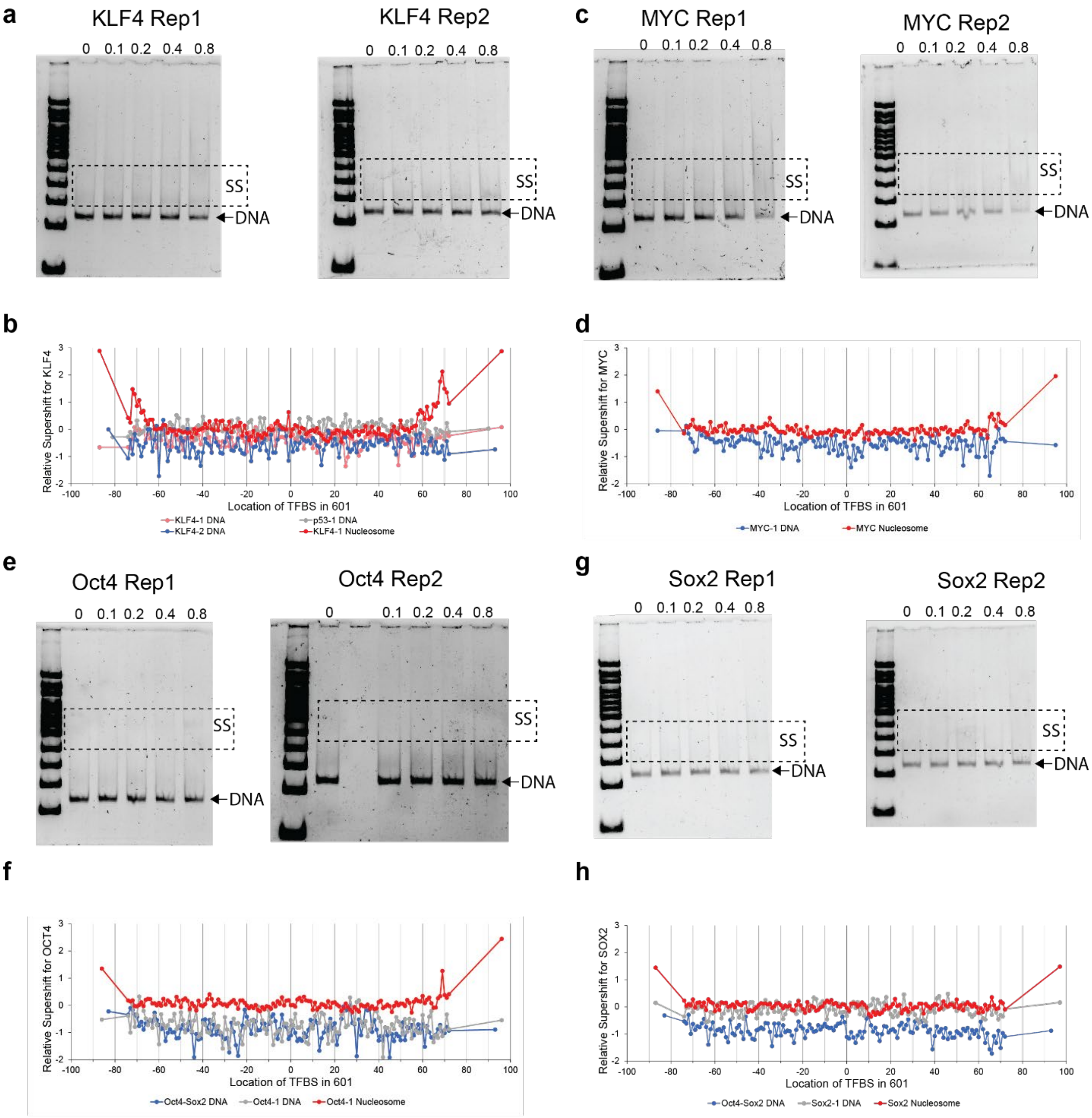


**Fig. S7. Naked DNA binding assays. (a,c,e,g)** 7500 sequences were bound to increasing amounts of TF and separated by native PAGE. All assay lanes contain 0.2 pmol DNA with 0, 0.1, 0.2, 0.4, or 0.8 pmols TF. Nucleosome band and the supershift (SS) region are indicated. (**b,d,f,h**) Relative supershift for the TF to naked Widom-601 DNA with the indicated TFBS. As a reference, the red line shows the relative supershift from Pioneer-seq to the indicated nucleosomes. (**b**) Relative supershift for 0.4 pmol KLF4. (**d**) Relative supershift for 0.8 pmol MYC/MAX. (**f**) Relative supershift for 0.2 pmol OCT4. (**h**) Relative supershift for 0.2 pmol Sox2.

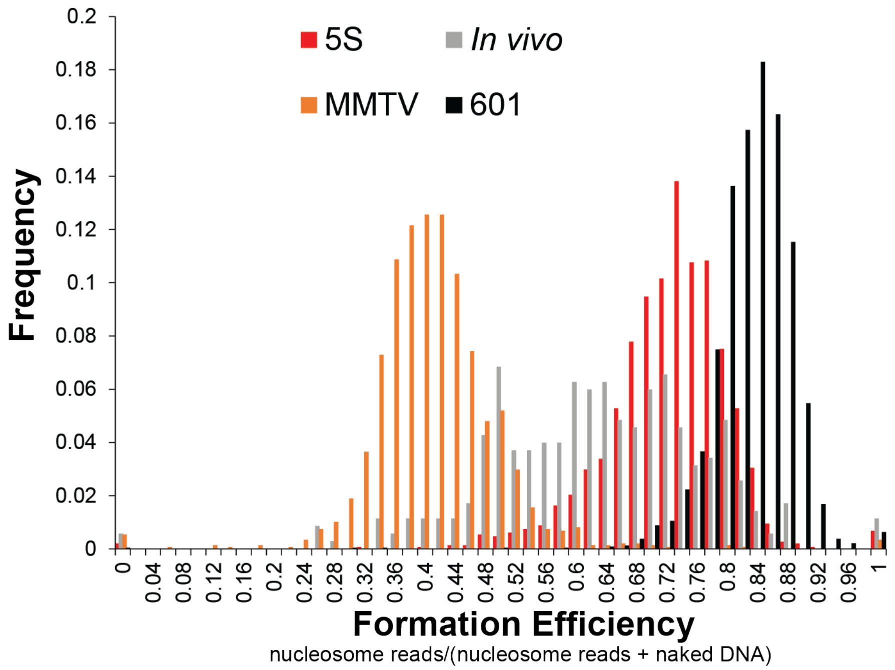


**Fig. S8. Nucleosome formation efficiency.** Nucleosome formation efficiency is determined before nucleosomes are purified from naked DNA by comparing the read numbers for every sequence in the 7500 library to the reads in the naked DNA band.

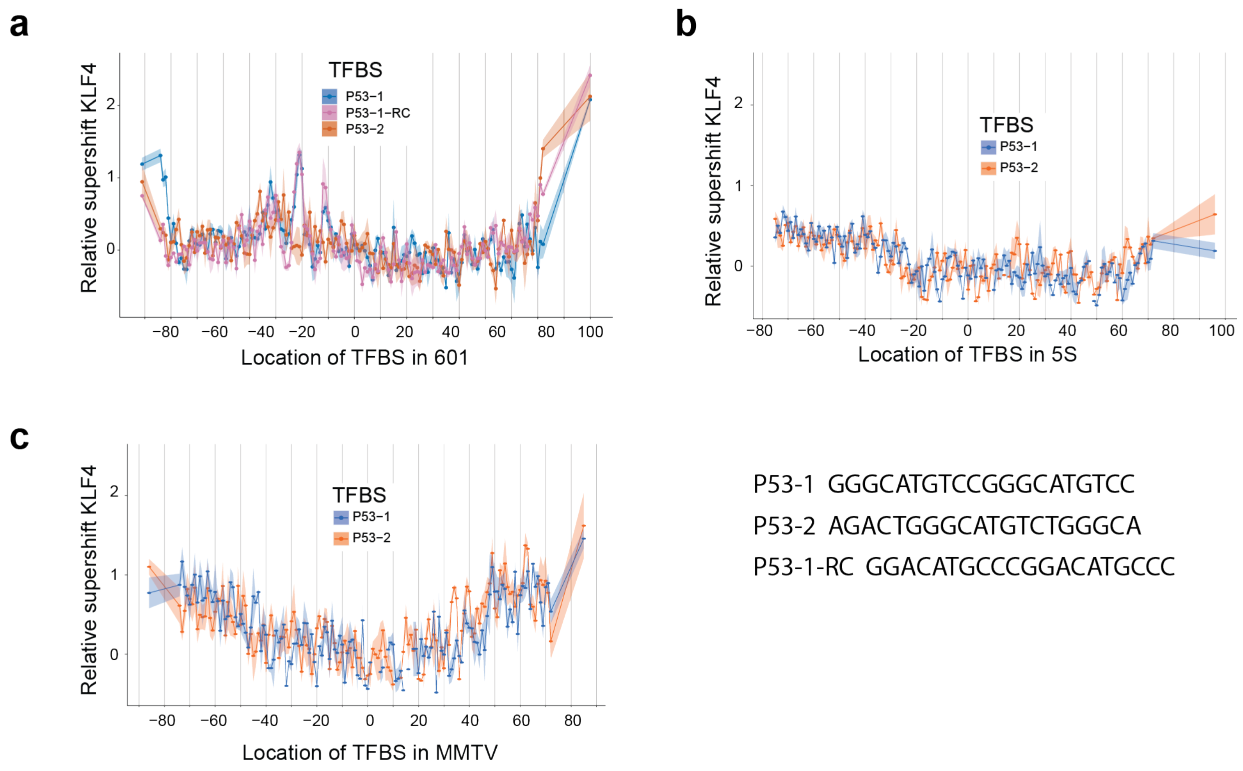


**Fig. S9. KLF4 binding to p53 binding sites.** The p53 TFBS are positioned across all possible locations along the (**a**) Widom-601 nucleosome, (**b**) 5S nucleosome, or (**c**) MMTV nucleosome with TFBSs in the left and right linkers to generate a total of 149 unique nucleosomes per TFBS. The relative supershift for each nucleosome is determined by counting the frequency of each sequence within the shifted band in the EMSA and comparing it to that for nonspecific binding. This value is then normalized to the input ratio of nucleosomes (see Eq. 1). Shading around each line is SEM.

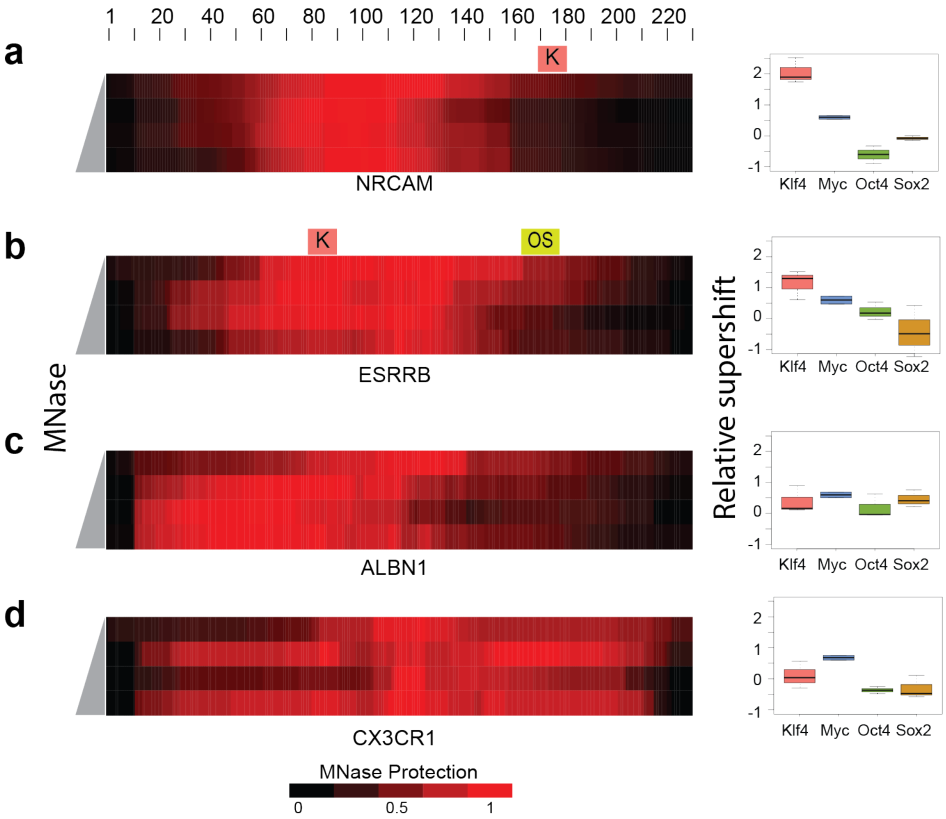


**Fig. S10. Binding to *in vivo*-nucleosomes from other studies.** The locations of TFBSs with MNase protection for *in vivo*-targeted nucleosomes (ITNs) are shown (red color scale at bottom). MNase protection was measured as the percentage of nucleosome bases that were protected from MNase digestion and calculated for each base pair as the ratio of base-pair coverage to the total reads for that specific nucleosome: **(a)**, NRCAM nucleosome from Garcia *et al* 2019. **(b)** ESRRB nucleosome from Huertas *et al* 2020. (**c)** ALBN1 nucleosome from Garcia *et al* 2019. (**d**) CX3CR1 nucleosome from Garcia *et al* 2019. The relative supershifts for each nucleosome are shown for KLF4, MYC, OCT4, and SOX2 binding on the right.

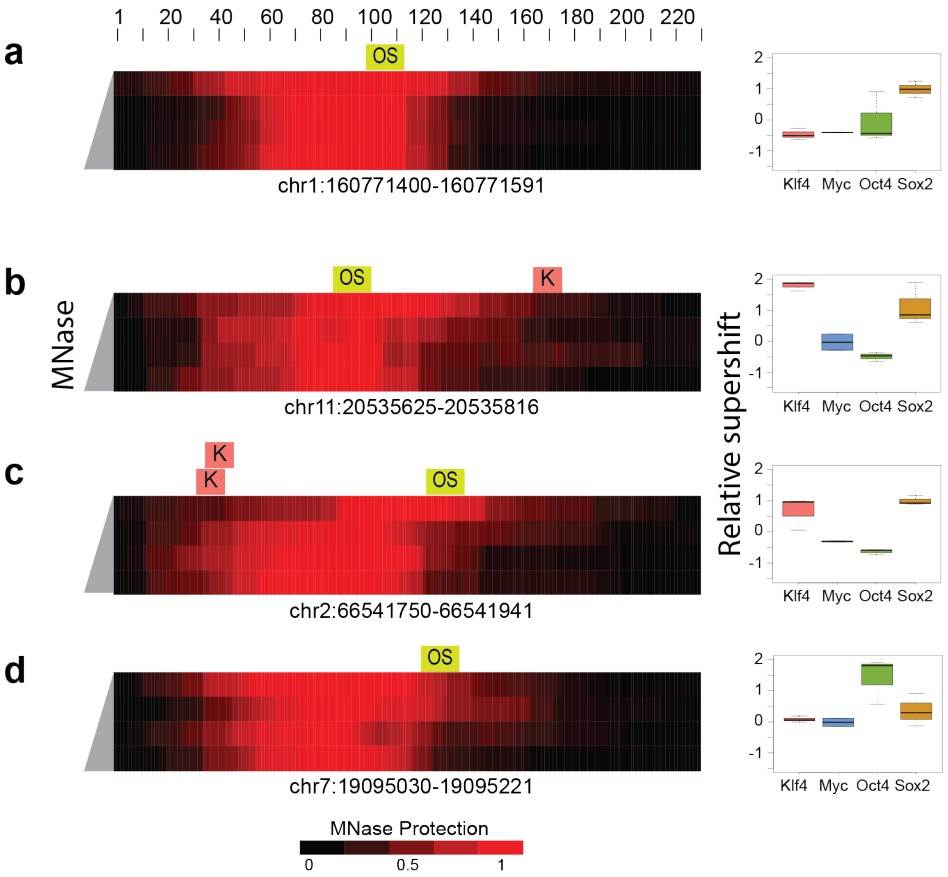


**Fig. S11. Binding to *in vivo*-targeted nucleosomes with Oct4-Sox2 binding site. (a-d)** The locations of TFBSs with MNase protection for *in vivo*-targeted nucleosomes (ITNs) are shown (red color scale at bottom). MNase protection was measured as the percentage of nucleosome bases that were protected from MNase digestion and calculated for each base pair as the ratio of base pair coverage to the total reads for that specific nucleosome.
